# Impact of climate change and oligotrophication on quality and quantity of lake primary production: A case study in Lake Biwa

**DOI:** 10.1101/2023.12.15.571786

**Authors:** Takehiro Kazama, Kazuhide Hayakawa, Takamaru Nagata, Koichi Shimotori, Akio Imai

## Abstract

Global climate change and anthropogenic oligotrophication are expected to reshape the dynamics of primary production (PP) in aquatic ecosystems; however, few studies have explored their long-term effects. In theory, the PP of phytoplankton in Lake Biwa may decline over decades due to warming, heightened stratification, and anthropogenic oligotrophication. Furthermore, the PP of large phytoplankton, which are inedible to zooplankton, along with biomass-specific productivity (PBc), could decrease. In this study, data from 1976–2021 and active fluorometry measurements taken in 2020 and 2021 were evaluated. Quantitatively, the temporal dynamics of mean seasonal PP during 1971–2021 were assessed according to the carbon fixation rate to investigate relationships among environmental factors. Qualitatively, phytoplankton biomass, PP, and PBc were measured in two size fractions [edible (S) or inedible (L) for zooplankton] in 2020 and 2021, and the L:S balance for these three measures was compared between 1992 (low-temperature/high-nutrient conditions) and 2020–2021 (high-temperature/low-nutrient conditions) to assess seasonal dynamics. The results indicated that climate change and anthropogenic oligotrophication over the past 50 years have diminished Lake Biwa’s PP since the 1990s, impacting the phenology of PP dynamics. However, the L:S balance in PP and PBc has exhibited minimal change since 1992. These findings suggest that, although climate change and oligotrophication may reduce overall PP, they do not markedly alter the inedible/edible phytoplankton balance in terms of PP and PBc. Instead, as total PP declines, the production of small edible phytoplankton decreases proportionally, potentially affecting trophic transfer efficiency and material cycling in Lake Biwa.

## Introduction

Globally, climate change is impacting organisms and transforming their ecosystems, including phytoplankton, a vital contributor to aquatic production and CO_2_ absorption (Finkel et al., 2010; Van de Waal & Litchman, 2020; Raven & Beardall, 2021). Phytoplankton production, i.e., primary production (PP), an indicator of lake and ocean productivity, is affected by climate change, yet accurately predicting the long-term effects at specific locations is challenging. For example, although rising sea surface temperatures are expected to decrease global PP (Steinacher et al., 2010), Arctic PP may increase (Frey et al., 2020). However, a decade-long mesocosm warming experiment did not show altered system PP (Padfield et al., 2018). Geographical and ecosystem-scale differences, along with long-term changes in light conditions (Ostrovsky et al., 2012) and nutrient loading (Koschel et al., 2002; Verbeek et al., 2018), further complicate predictions in natural environments. Regarding the latter, its global implementation in terrestrial waters has garnered attention for its interplay with climate change (Verbeek et al., 2018). Despite this, previous studies assessing long-term shifts in lake ecology, including paleontology, have been limited to discussions spanning annual to decadal scales (Tsugeki et al., 2010; Tsugeki & Urabe, 2012).

Climate change not only affects the quantity of PP but also its quality. Strengthened stratification and oligotrophic conditions prevail in the mixed layer under global warming, favoring small phytoplankton dominance (Falkowski & Oliver, 2007; Winder et al., 2009; Zohary et al., 2021). Given that key herbivorous zooplankton, such as *Daphnia* and copepods, feed on smaller PP sizes, with 20–50 μm being the limit (Lampert & Sommer, 2007), downsizing PP affects trophic structure (Kazama et al., 2021c) and material cycling (Ray et al., 2001; Law et al., 2009). Therefore, considering long-term PP changes necessitates examining the qualitative aspect of size structure.

The balance between large inedible phytoplankton (L) and small edible phytoplankton (S) production is a crucial consideration. PP is determined by multiplying carbon-specific productivity (PBc) by carbon biomass; therefore, PP changes with alterations in the L:S biomass balance [refer to reviews on the determination mechanism for the L:S balance of biomass by Zohary et al. (2021) and Hillebrand et al. (2022)]. However, the long-term changes in the PBc of natural phytoplankton communities have not been reported. Although the PBc of S is typically larger than that of L (Banse, 1976; Finkel et al., 2010; Marañón, 2015; Hillebrand et al., 2022), and it is assumed that the PBc L:S balance remains stable in natural environments, instances of PBc with L > S have been reported in natural communities (Cermeño et al., 2005b; Liu et al., 2021), suggesting variable PBc L:S balances. Such cases are often reported in marine coastal zones, characterized by favorable conditions for large phytoplankton owing to high vertical mixing, nutrient inputs, and abundant light (Iriarte & Purdie, 1994; Maraóón et al., 2007; Li et al., 2020). Consequently, it is plausible that stratification enhancement due to warming or reduction in nutrient loading could result in a sustained decrease in the PBc L:S balance. Nevertheless, continuous data on long-term changes in PP and PBc L:S balance are lacking in oceans and lakes.

In Lake Biwa, surface temperature has increased by 0.357°C per decade since the 1970s (Fig. 1A, and winter vertical mixing has been reduced due to strengthened stratification (Yamada et al., 2021; Zhou et al., 2022). However, total nitrogen (TN) and total phosphorus (TP) concentrations have been reduced through restoration efforts since the 1970s, making Lake Biwa highly oligotrophic (Fig. 1D, E). Fish species and abundance under these conditions are lower relative to the 1960s (Liu et al., 2019; Nishino, 2020). Although these changes may be attributed to quantitative shifts in PP, it was only measured intermittently up to 2008 (Goto et al., 2014), with recent dynamics remaining unknown. After the first size-fractionated PP and PBc measurements in 1992 (Urabe et al., 1995), the large alga *Micrasterias hardyi* became more abundant and is now dominant with another large green alga, *Staurastrum dorsidentiferum* var. *ornatum* (hereafter, *S. dorsidentiferum*) (Hodoki et al., 2020; Kazama et al., 2022). Therefore, obtaining present size-fractionated PP and PBc data is crucial for evaluating long-term qualitative changes and quantitative changes over 50 years.

**Fig. 1.**
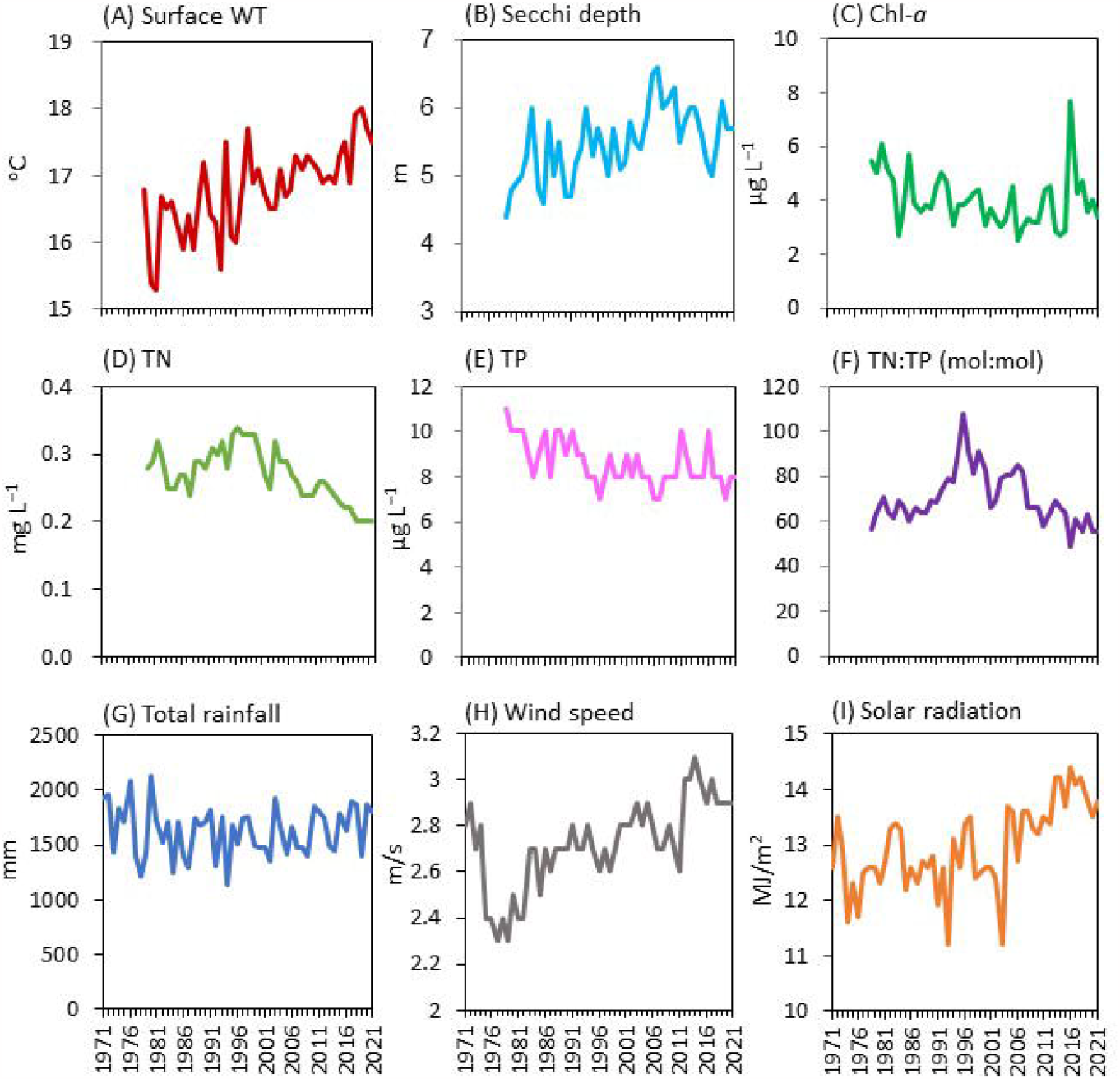
Annual changes in climate and water environments over 50 years in the North Basin of Lake Biwa. Water environments data were collected from Annual White Paper on the environment of Shiga Prefecture 2022 (Shiga Prefecture 2022). Climate data was collected from the Japan Meteorological Agency.

This study combines a literature review and field study to assess the long-term impacts of climate change and anthropogenic oligotrophication on the quantitative and qualitative aspects of PP. Quantitatively, the temporal dynamics of mean seasonal PP during 1971–2021 were evaluated based on carbon fixation rates to explore relationships with environmental factors. Qualitatively, phytoplankton biomass, PP, and PBc were measured in two size fractions [edible (S) or inedible (L) for zooplankton] in 2020 and 2021. Comparing seasonal dynamics of the L:S balance for phytoplankton biomass, PP, and PBc between 1992 (low-temperature/high-nutrient conditions) and 2020–2021 (high-temperature/low-nutrient conditions), the relationships among PP and PBc L:S balance and physical, chemical, and biological environments were analyzed to determine the mechanisms underlying PP and PBc L:S balance.

## Materials and methods

### Sampling

This study was conducted at two sampling stations established in Lake Biwa on Honshu Island, Japan (Fig. 1). Monthly sampling was conducted at the long-term survey stations, Station 12B (62 m depth, 35°11′39″ N, 135°59′39″ E) and Station 12C (7 m depth, 35°10′40″ N, 136°03′07″ E) in the North Basin, from April 2020 to December 2021. No permits were required for lake sampling in this study. Vertical profiles of irradiance and water temperature were measured using a water quality sonde (AAQ-RINKO; JFE Advantech Co., Ltd., Hyogo, Japan). Water samples were collected with 5 L Niskin bottles on a rosette sampler (AWS; JFE Advantech Co. Ltd., Kobe, Japan) at 5.0 m at Station 12B and with a 10 L Niskin bottle at 2.5 m at Station 12C. Samples were then stored in plastic bags in the dark and transferred to the laboratory for chemical and biological analyses within 2 h after collection.

Sampled water was initially filtered through 20 μm nylon mesh to eliminate crustacean zooplankton (major grazers in Lake Biwa) and then separated into two size classes, small (S < 30 μm) and large (L > 30 μm), using 30 μm nylon mesh. Given the observation of aggregated cyanobacterial colonies, such as *Aphanocapsa, Aphanothece*, and *Chroococcus*, in the L class in August 2020 (Appendix Fig. 1), a 45 μm nylon mesh was used for size fractionation in 2021. However, these aggregated colony were not excluded. Given that dominant species of large phytoplankton, such as *S. dorsidentiferum* and *M. hardyi*, were larger than 45 μm, the separation of phytoplankton communities using 30 or 45 μm mesh yielded similar results. Size-fractionated chlorophyll a (Chl-a) and macronutrient concentrations were analyzed using a previously established method (Kazama et al., 2022). For seston C, N, and P, 50–300 mL size-fractionated samples were filtered with precombusted 25 mm glass fiber filters (0.7 μm nominal pore size; GF/F; GE Healthcare, UK Inc., Little Chalfont, UK). Seston samples were dried at 56°C for 24 h and stored in a desiccator until analyses. Seston C and N were analyzed using an elemental analyzer (Flash EA 1112; Thermo Finnigan, Bremen, Germany). Seston P was analyzed using a continuous flow analyzer (QuAAtro 2-HR; BLTEC, Tokyo, Japan) following potassium persulfate digestion.

To determine phytoplankton numbers, a 50 mL aliquot of each sample was fixed with Lugol’s solution (1% final concentration). After settling in darkness for 24 h, the supernatant was gently removed, and the sample was concentrated to 15 mL. All phytoplankton cells were enumerated at the finest taxon level (species or genus) under a light microscope at ×100–400, with biomass estimated according to cell volume following an established methodology (Kazama et al., 2021a, 2022).

### Size-dependent photophysiology

Preliminary screened samples were fractionated into the two size classes mentioned earlier. Large phytoplankton were resuspended in lake water filtered through a GF/F filter, and diluted to the *in-situ0020*concentration. Photophysiology was measured using a previously described method (Kazama et al., 2021b, 2022). Briefly, 2 mL samples from each size class were poured into two 10 mL glass tubes. After 20 min acclimation to low irradiance (1–10 μmol m^−2^ s^−1^), maximum photochemical efficiency (*F*_*v*_*/F*_*m*_), effective photochemical efficiency under actinic light (*F*_*q*_*′/F*_*m*_*′*) as PSII photochemistry efficiency, and PSII light absorption cross-section [σ _PSII_(BGO)] were measured using blue, green, and orange flashlets from the bench-top FRRf (Act2, CTG Ltd., West Molesey, UK). Data with *Rσ*_*PSII*_ (the probability of an RCII being closed during the first flashlet of a single turnover saturation phase under dark conditions) or *Rσ*_*PSII*_*′* (similar to *Rσ*_*PSII*_ but under actinic light) of <0.03 or >0.08 were excluded from further analysis.

### Size-dependent productivity

The electron transport rate per water volume (*JV*_*f*_, μmol electrons m^−3^ s^−1^) was derived following Kazama et al. (2021a). To address differences in spectral distribution and primary production response between the excitation flash and actinic light sources in the Act2 system and ambient light in water column, a spectral correction factor (SCF) was applied to the dataset to mitigate potential discrepancies between the methods (Schuback et al., 2017). The SCF was calculated using the following formula:

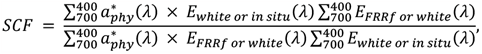

where 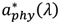 is the Chl-*a–*specific absorption spectrum of phytoplankton (m^2^ mg Chl-*a*^−1^), and *E*_*white*_ (λ), *E*_*in situ*_ (λ), and *E*_*FRRf*_ (λ) are the spectral distributions of the actinic light, water column, and excitation flash, respectively. 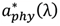 was measured using a quantitative filter technique (IOCCG, 2018) employing Lambda 850+ (PerkinElmer Inc., Waltham, MA, USA) with an integrating sphere. *E*_*in situ*_ (λ) was measured using a hyperspectral irradiance sensor (Ramses; TriOS Mess- und Datentechnik GmbH, Germany).

The primary productivity per water volume (PP mg C m^−3^ h^−1^) was determined through *JV*_*f*_ with an electron requirement for carbon fixation (Ф_e,C_) using the following formula:

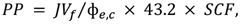

where 43.2 is the conversion factor from seconds to hours and from μmol C to mg C. Ф_e,C_ was derived from an empirical model based on the ^13^C tracer method, including physical and chemical factors, photosynthetic parameters, and taxonomic composition as explanatory variables (Kazama et al., 2021a).

PB_C_ (mg C mg C^−1^ h^−1^) was calculated from PP/seston C. PBmax and α for PB_C_ were derived from a production–light curve fitted in a two-parameter model, as described by Webb et al. (1974).

## Data collection

Climate data were collected from the Japan Meteorological Agency. Lake environmental data were obtained from Annual White Paper on the Environment of Shiga Prefecture 2022 (Shiga Prefecture, 2022). PP data were compiled from previous measurements based on the ^14^C or ^13^C method (Nakanishi, 1976, 1984; Nakanishi et al., 1992; Tsuda & Nakanishi, 1992; Takahashi et al., 1995; Urabe et al., 1995, 2005; Goto et al., 2014; Kazama et al., 2021a), as O_2_ production rates do not necessarily equate to carbon fixation rates (Halsey et al., 2010; Regaudie-de-Gioux et al., 2014). In a previous study, PP values determined via a longer (>4h) incubation time were considered rather net PP (Kromkamp et al., 2017); thus, we present these data (1983–1984 and 1992) with notifications (Fig. 2), and excluded these from regression analysis (Fig. 3).

**Fig. 2.**
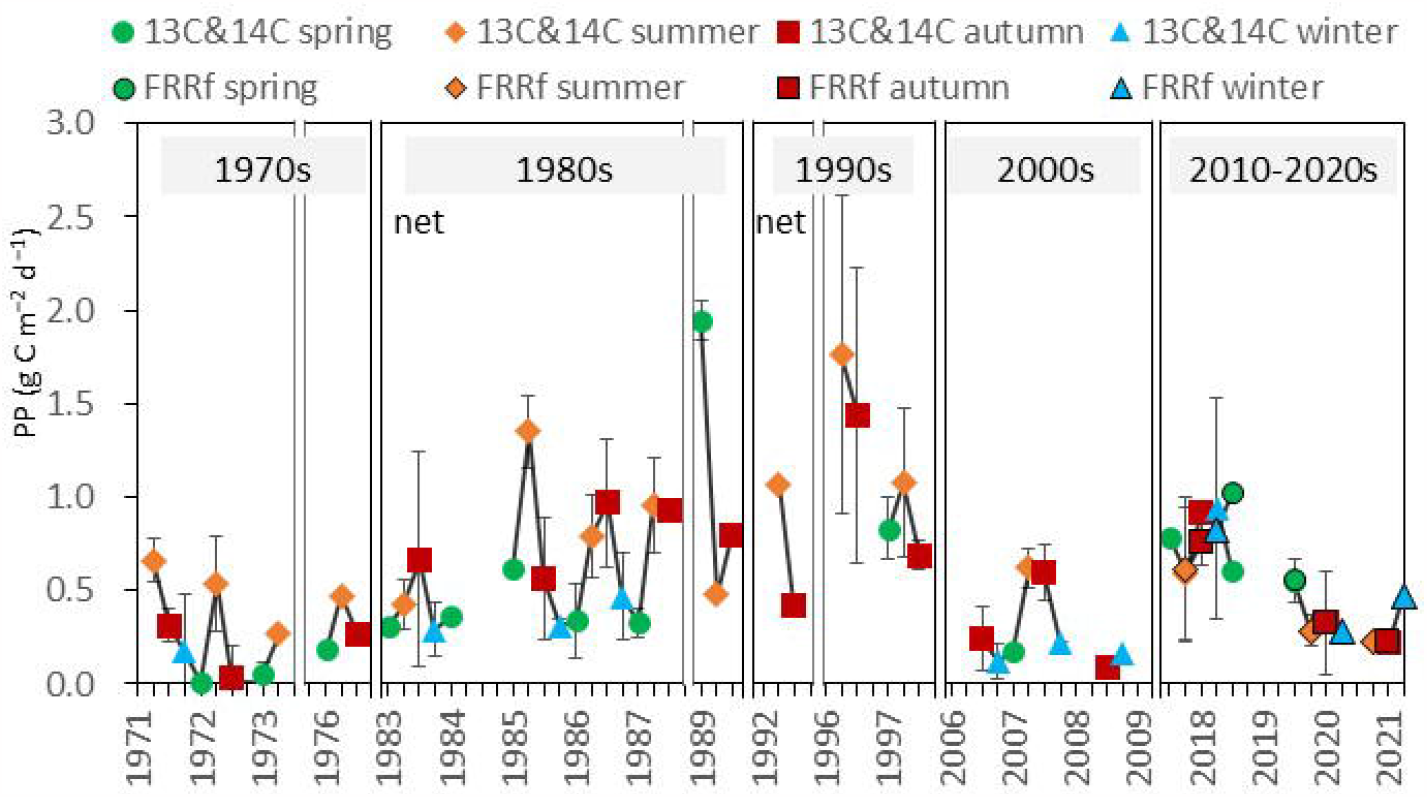
Seasonal average of primary productivity per area (PP) in the North Basin of Lake Biwa in spring (March– May), summer (June–August), autumn (September–November), and winter (December–February) from 1970s to 2020s. Values were compiled from previous measurements based on the ^14^C or ^13^C method in previous studies (**1970s**: Nakanishi 1976; Nakanishi 1984. **1980s**: Tsuda and Nakanishi 1992; Takahashi et al. 1995; Nakanishi et al. 1992. **1990s**: Urabe et al. 1995; 2005. **2000s**: Goto et al. 2014. **2010-2020s**: Kazama et al. 2021; This study). Solid vertical lines denote border of the decades. Dotted vertical lines denote border of the study periods within a decade. PPs determined by longer (> 4h) incubation time are indicated as “net” (see “Data collection”). Error bar denotes standard deviation.

**Fig. 3.**
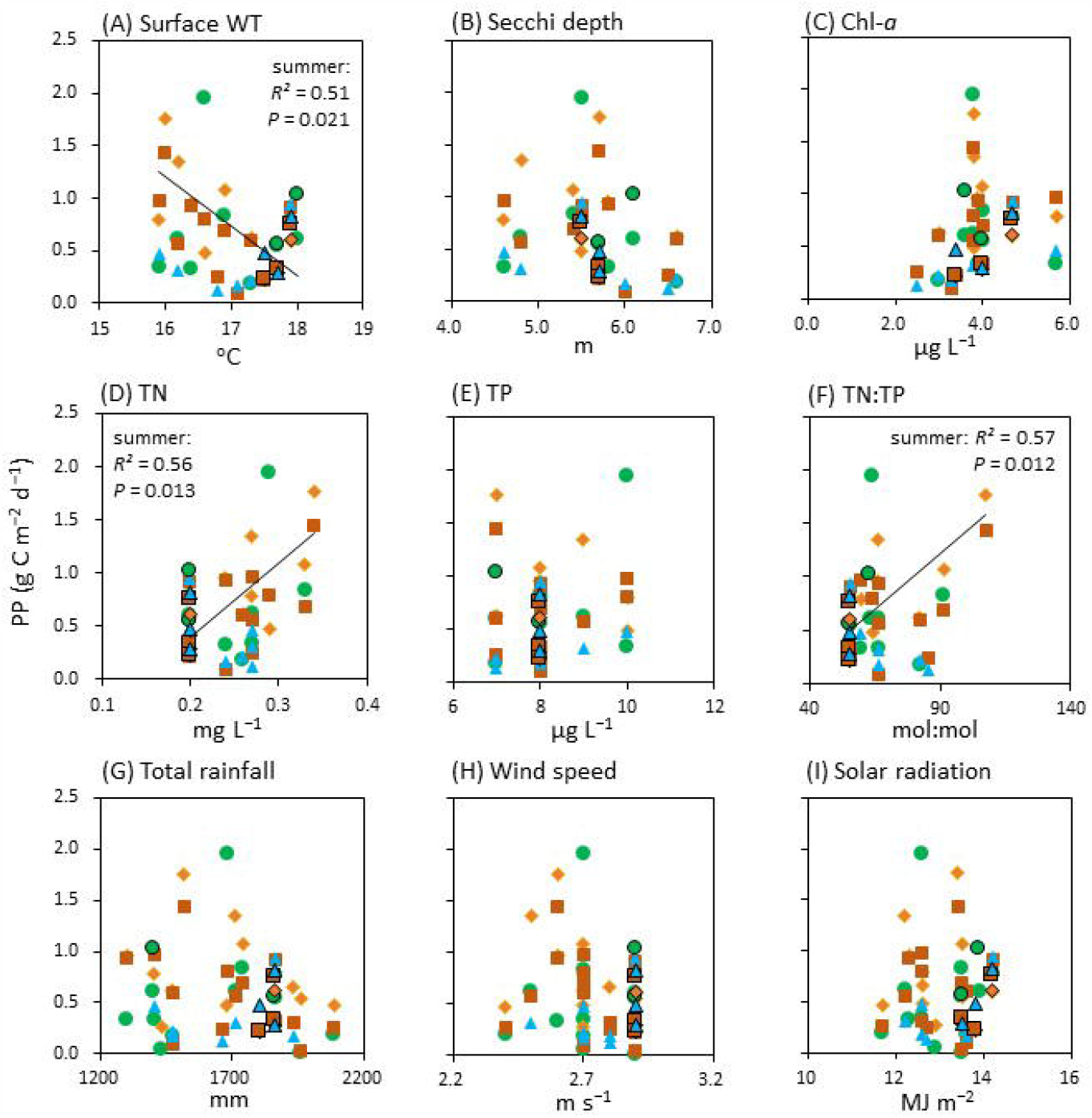
Relationships between PP and average surface water temperature (A), secchi depth (B), Chl-a (C), TN (D), TP (E), TN:TP ratio (F), total rainfall (G), average wind speed (H), and average solar radiation (I). Data from 1983– 1984 and 1992 were excluded because these values are rather net production (see “Data collection”). *R*^*2*^ and *P*-values were estimated by Pearson’s correlation test.

The L:S balance in seston C, PP, and PBc in the North Basin of Lake Biwa in 1992 were calculated using the data of Urabe et al. (1995). Given that PP data from 1992 were considered net PP, as mentioned earlier, we focused on the L:S balance. Notably, the community separation procedure used by Urabe et al. (1995), filtering algal communities using 20 μm mesh, differed from our chosen procedure. However, *S. dorsidentiferum* and *Closterium aciculare* var. *subpronum*, the dominant species in 1992 (The Shiga Prefectural Institute of Public Health and Environmental Science, 1995), do not pass through the 30 and 45 μm mesh used in our study. Nevertheless, it is our contention that filtration mesh size should not influence the community size spectrum, and the L:S balances of seston C, PP, and PBc can, therefore, be compared between the 1992 and 2020–2021 datasets.

### Statistical analyses

Pearson’s correlation test, performed in R.4.3.0. (R Development Core Team, 2023), was used to test relationships between environmental factors and PP. Changes in seston C, and PP, and PBc L:S balances between 1992 and 2020 or 2021 were tested in R using the Steel test via the nparcomp() function in the “nparcomp” package (Konietschke et al., 2015). For comparisons between 1992 and 2020–2021, June–November data from each sampling site as well as the data of Urabe et al. (1995) were used. Differences in PP, PBc, and photosynthetic parameters between L and S during April 2020 and December 2021 were also tested using the Wilcoxon signed-rank test.

Redundancy analysis (RDA) was conducted to explore the relationships between the L:S balance of photosynthetic parameters and physical and chemical environments, along with taxonomic composition. To align with normality assumptions, data underwent log transformation before RDA application. For log transformation of taxonomic composition data, log(x + 0.01) was applied to each group. RDA was performed using the rda() function in the “vegan” package (Oksanen et al., 2022) in R. The statistical significance of each component in RDA was evaluated through permutation-based ANOVA (1000 permutations). Both the code and data used in this study are available in Zenodo (https://doi.org/10.5281/zenodo.10376510).

## Results

### Climate, water environment, and PP changes over 50 years

Over the last 50 years, the lake’s water environment has exhibited rising temperatures (Fig. 1A) and increased transparency (Fig. 1B). Chl-a content has decreased, with an exception in 2016 (Fig. 1C), and TN and TP have decreased since the late 1990s (Fig. 1D, E). However, TP stabilized over the following two decades, and the TN:TP peaked in 1996 before declining to reach its lowest level in 50 years (Fig. 1F). The climate surrounding Lake Biwa has undergone changes, with increased wind speed (Fig. 1H) and improved light conditions (Fig. 1I) reported since the late 1970s.

Figure 2 shows the seasonal average of PP for each decade from 1971 in spring (March–May), summer (June–August), autumn (September–November), and winter (December–February). Although PP typically peaked in summer or autumn from the 1970s to the 2000s, its peak shifted to spring and winter during the 2010s to 2020s. Additionally, seasonal variation in PP was minimal over these two decades. Despite potential errors in the 2020 and 2021 empirical Ф_e,C_ estimation, PP values measured via the ^13^C tracer method reflected a similar trend during 2018–2019.

The correlation between seasonal PP and annual averages of climate and water environments (Fig. 1) indicated a significant decrease in summer PP with rising water temperature (*R*^*2*^ = 0.51; *p* = 0.021; Fig. 3A). Conversely, summer PP increased significantly with elevated TN (*R*^*2*^ = 0.56; *p* = 0.013; Fig. 3D) and TN:TP (*R*^*2*^ = 0.57; *p* = 0.012;Fig. 3IF. In recent years, the lowest summer PP values were associated with high water temperature and low TN (Fig. 4).

**Fig. 4.**
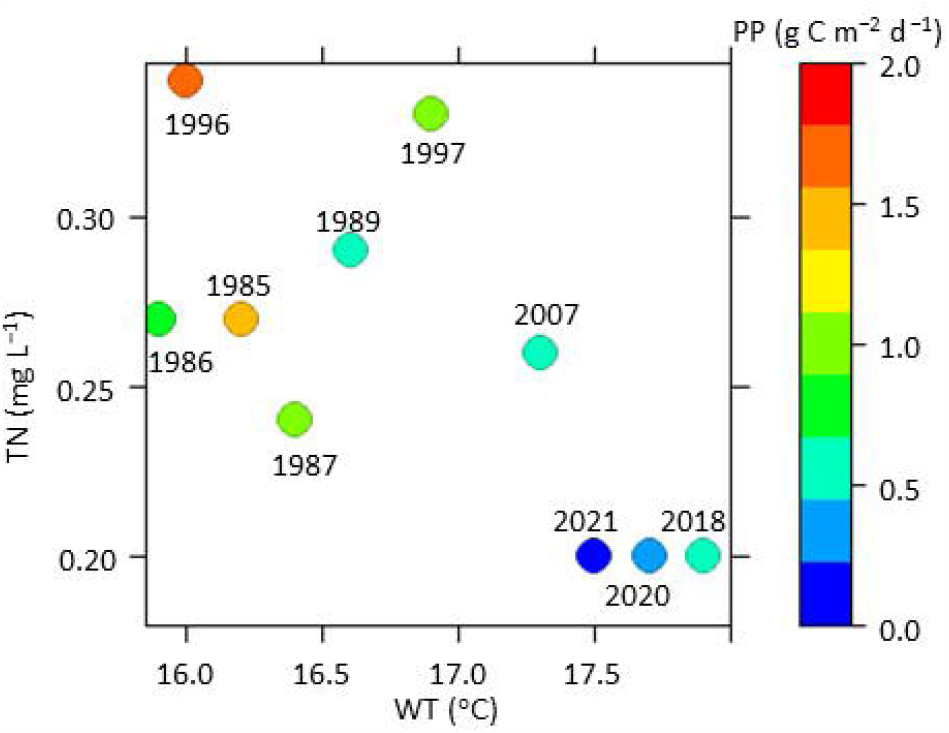
Summer PP against water temperature (x-axis) and TN (y-axis).

### L:S balance of seston C, PP, and PBc in 1992, 2020, and 2021

Comparisons of the seasonal dynamics of seston C, PP, and PBc L:S balance between the low-temperature/high-TN year, 1992, and the high-temperature/low-TN years, 2020–2021, are shown in Fig. 5 (refer to Appendix Fig. 2 for comparisons of absolute values). In 1992, the seston C L:S balance was around 1 in June, decreased to <1 in August and September, and increased again in November (Fig. 5A). In 2020 and 2021, although similar patterns were observed in Station 12B and Station 12C, values at Station 12B were mostly higher than those at Station 12C. A significant decrease in the mean seston C L:S balance was observed in 2021 from June to November compared with 1992 (Steel test, *P* < 0.05; Fig. 5B; Appendix Table 1).

**Fig. 5.**
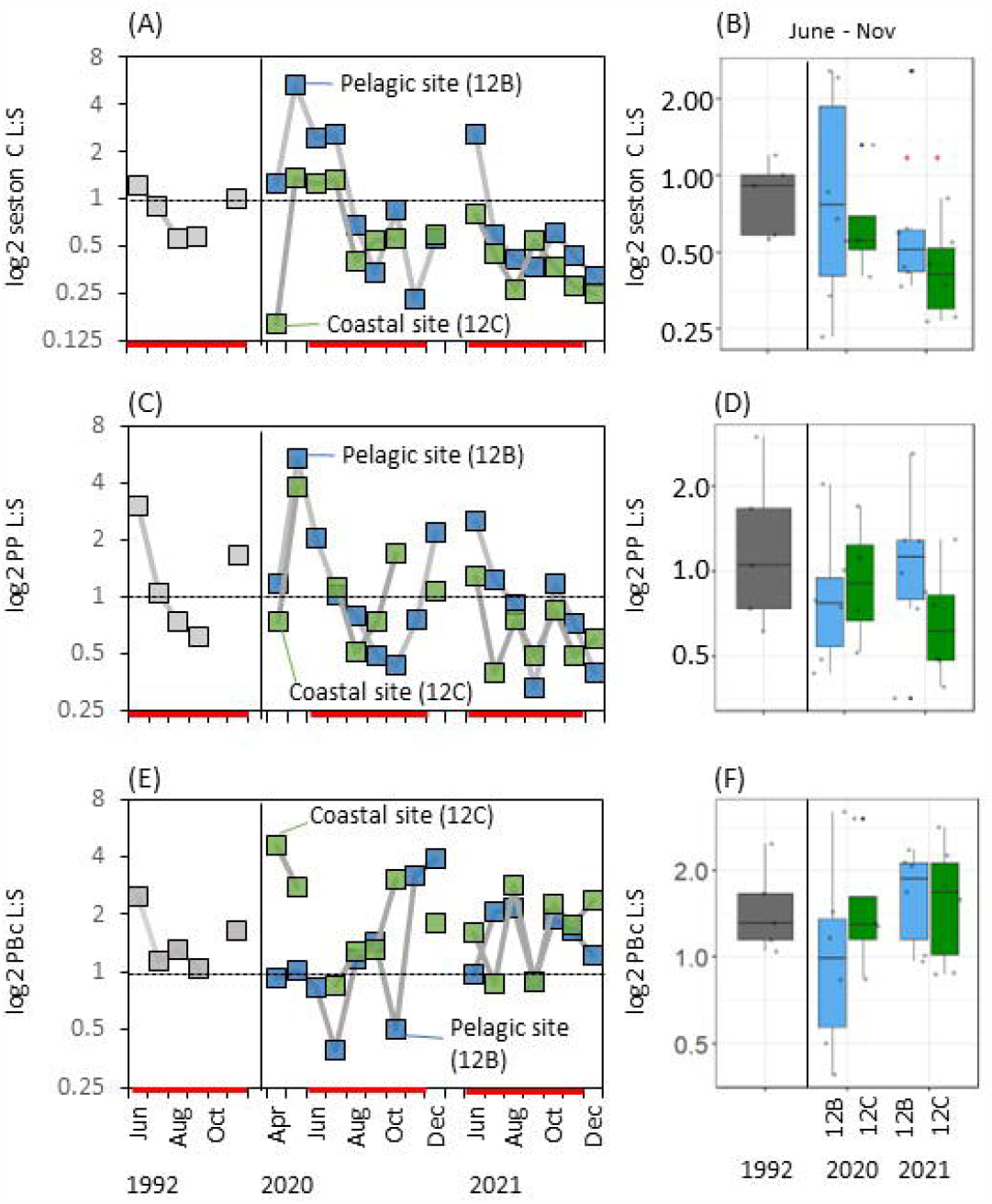
**(A, C, E):** Temporal changes in the log2 L : S balance in seston C, PP and PBc in the North Basin in Lake Biwa in 1992 (Urabe et al. 1995) and 2020–2021 (This study). Vertical line denotes border of the decade. Horizontal dashed line denotes L : S = 1 : 1. Periods used for comparisons in the right panels were highlighted by bold red lines. **(B, D, F):** The box plots for each L : S balance from June to November. The whiskers indicate 1.5 times the interquartile range (Q3−Q1) below and above Q1 and Q3. Outliers beyond the whiskers were plotted individually. *, *P* <0.05 for comparison between 1992 and 2020 or 2021 for 12B or 12C by Steel’s test.

The PP L:S balance exhibited a comparable seasonal pattern in seston C (Fig. 5C), peaking at >1 in early summer and then falling to <1 in late summer and early autumn. This pattern aligned with the seasonal dynamics of *S. dorsidentiferum* and *M. hardyi* (Appendix Fig. 3). No significant differences were found between 1992 and 2020 or 2021 in the mean PP L:S balance from June to November (Fig. 5D; Appendix Table 1).

The PBc L:S balance remained mostly above 1 in 1992 and at Station 12C in 2020–2021, except for a drop below 0.5 at Station 12B in July 2020 (Fig. 5E). However, no significant differences in this balance were observed between 1992 and 2020 or 2021 from June to November (Fig. 5F; Appendix Table 1).

### Size-fractionated photosynthetic parameters, PBc, and PP in 2020–2021

Comparisons of photosynthetic parameters [*F*_*v*_*/F*_*m*_ and σPSII(BGO)], PP, PBc, and PB parameters (PBc max and αc) for April–December 2020 and June–December 2021 are shown for each site in Fig. 6. Despite using different mesh sizes for fractionation in 2020 and 2021, a Wilcoxon test was applied to pooled data because the dominant species *S. dorsidentiferum* and *M. hardyi*, both larger than 45 μm, showed similar PP and PBc variations in 2020 and 2021 (Appendix Fig. 2C, E). Although PP did not differ significantly between sizes (Fig. 6A), *F*_*v*_*/F*_*m*_ was higher in fraction L, whereas σ_PSII_(BGO) was higher in fraction S (Fig. 6B, C) at Station 12B and Station 12C. Furthermore, PBc, PBc max, and α c were higher in fraction L at Station 12C, whereas no significant differences were observed at Station 12B (Fig. 6D–F).

**Fig. 6.**
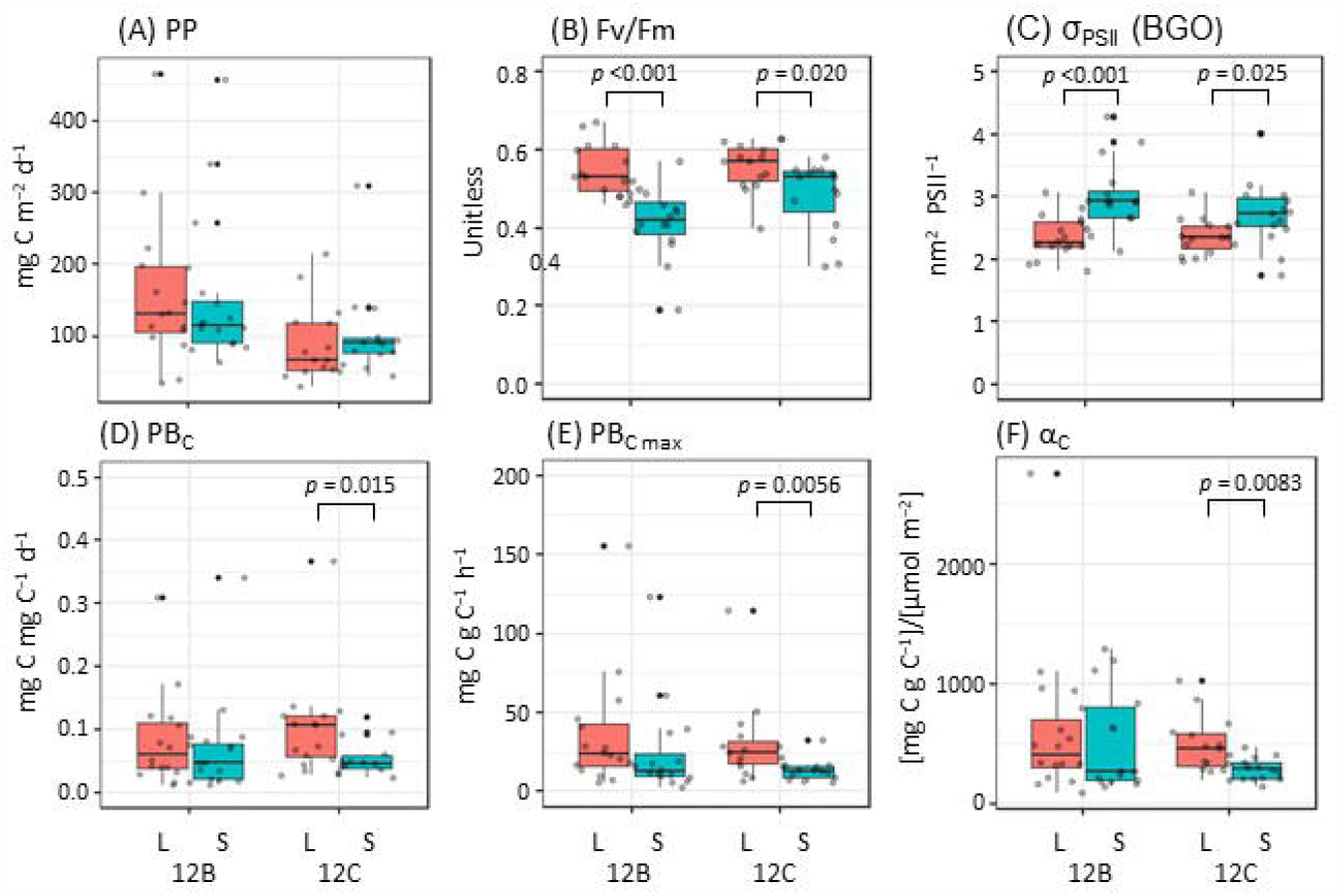
Box plots for photosynthetic and PB-parameters in pelagic (12B) and coastal (12C) sites during 2020–2021. Median, Q1, Q3, whiskers and outliers were shown. Significant (*P* < 0.05) differences between L and S are shown with *P*-values of Wilcoxon signed rank test in each site.

### Relationships between environmental factors and PB parameters

The relationships between the L:S balance of PB parameters (PBc max, αc, PBc, and PP) and environmental variables were examined via RDA (Fig. 7). The first and second axes of the RDA explained 52.0% and 24.4% of the variation in PB parameters, respectively. Through forward selection, four significant environmental variables were found to explain spatial and temporal changes in PB parameters. These variables included the Chl-a L:S balance (chl.aLS, *F* = 10.28, *P* = 0.003), seston C L:S balance (sestonCLS, *F* = 4.09, *P* = 0.031), Chl-a:seston C ratio L:S balance (chl.CLS, *F* = 8.38, *P* = 0.030), and 4-day average irradiance before the sampling day (I.4ave, *F* = 4.66, *P* = 0.020). RDA results revealed that PBc, PBc max, and αc were positively correlated with the Chl-a:seston C ratio L:S balance, whereas the PP L:S balance was positively correlated with the Chl-a and seston C L:S balances, along with the 4-day average irradiance.

**Fig. 7.**
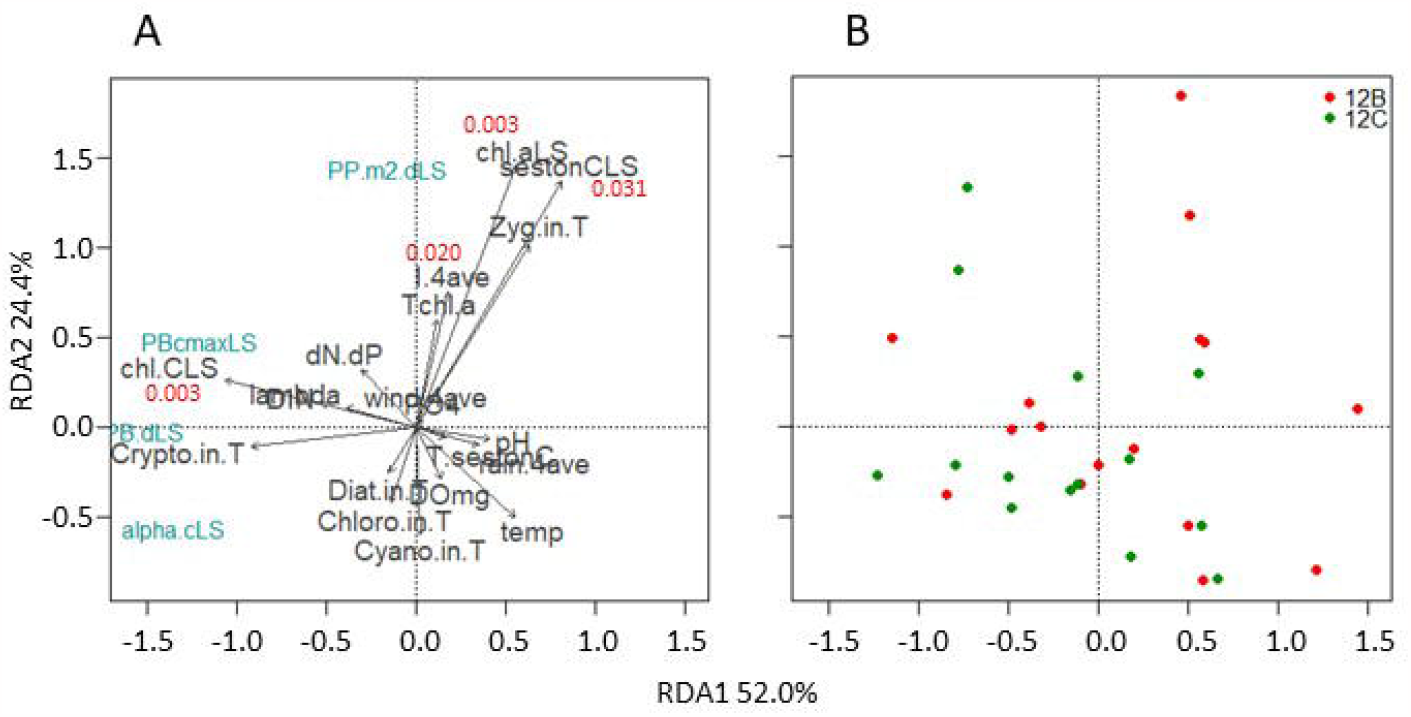
Biplots of redundancy analysis (RDA) for L:S balane of PB-parameters (PBc max, αc, PBc and PP) and environmental variables (A), and site scores (B). The first two axes explain 76.3% of variation in photosynthetic parameters, and 99.3% of variation in environment data, respectively. The goodness of fit of each parameters to the presented ordination is represented by the arrow length. The degree of correlation is shown as the angle between arrows. Abbreviations of the environmental variables are: temp, water temperature; chl.C, Chl-a:seston C ratio; Diat-/Chloro-/Cyano-/Crypto-/Zyg .in.T, diatoms-/chlorophytes other than zygnematophytes-/cyanobacteria-/cryptophytes-/zygnematophytes in total biomass; I-/rain-/wind. 4ave, 4-days average of irradiance/rainfall/windspeed before sampling day. Significant environmental variables (*P* <0.05) are denoted with *P-*values in red.

## Discussion

### PP dynamics over 50 years

The trends in PP dynamics from the 1960s to the 2000s closely align with the ^13^C stable isotope ratios in sediments (Hyodo et al., 2008), suggesting that past PP measurements are representative of each decade. Notably, PP has been consistently decreasing since the late 1980s. The negative relationship between summer PP and physical and chemical environments implies the direct and indirect effects of warming and oligotrophication. For example, Garcia-Corral et al. (2017) found a negative correlation between PP in subtropical and tropical oceans and water temperature in the range 16°C–29°C. Additionally, our previous study indicated a decrease in carbon fixation efficiency with rising temperatures (Kazama et al., 2021a). Increased water stratification and a reduced mixed layer may diminish nutrient supply to the surface layer, leading to decreased ocean PP (Steinacher et al., 2010). Given Lake Biwa’s deep and oligotrophic nature, weakened vertical mixing (Yamada et al., 2021) may induce nutrient limitation in the upper layer.

Anthropogenic oligotrophication is expected to be a major factor, independently or in conjunction with warming, contributing to the decline in PP. Nutrient load reduction efforts, ongoing since 1970 in Lake Biwa (Nishino, 2012), have contributed to a decrease of approximately 0.1 mg L^−1^ and 2 μg L^−1^ in TN and TP, respectively, since the earliest record in 1979 (Fig. 1D, E). The demand for nitrogen, crucial for Chl-a synthesis in phytoplankton (Sterner & Elser, 2002), may increase with warming (Thomas et al., 2017), potentially exacerbating the negative effects of nitrogen limitation, particularly in summer. Jiang & Nakano (2022) suggested that, given current TN (0.20–0.33 mg L^−1^) and TP (7–11 μg L^−1^) concentrations, Lake Biwa’s nutrient environment is near the threshold for phytoplankton nitrogen limitation.

The seasonal PP pattern in the 2010s to 2020s deviated from the typical summer–autumn peak observed in the 1970s to 2000s, displaying a distinctive pattern with relatively high winter PP (Fig. 2). Although warming may negatively impact summer PP, the warmed mixed layer in winter could enhance PP in the photic layer zone, akin to effects observed in polar regions (Behrenfeld et al., 2006; Frey et al., 2020). Anticipated future increases in stratification (Donis et al., 2021) may further extend changes in PP phenology, potentially influencing the seasonal succession of zooplankton (Sommer et al., 2007).

### Changes in the L:S balance of PP and PBc

Comparing 1992 with 2020–2021, water temperature increased (1.2°C–1.4°C) while TN and TP decreased (0.1 mg L^−1^ and 1 μg L^−1^, respectively) in the later years. As predicted by Kishimoto et al. (2013), there was a significant decrease in the relative biomass of large phytoplankton, especially zygnematophytes, in 2021 (Fig. 5B; Appendix Fig. 4). Despite a slight decrease in the PP L:S balance at Station 12B in 2020 and Station 12C in 2021, both values fell within the range 0.5–1.5 (Fig. 5D). This is attributed to the relatively high PBc L:S balance counteracting the influence of decreased seston C L:S balance (Fig. 5F). Even considering the estimation error in Ф_e,C_ (maximum 2-fold; Kazama et al., 2021a), the PBc of large phytoplankton remains high, as observed in 1992. Additionally, the phenology of the PBc L:S balance showed no difference in 2020–2021. These results suggest that, despite a decline in the abundance of large phytoplankton, their productivity remains higher than that of small phytoplankton. Tilstone et al. (2017) evaluated 1998–2011 PP size fractionation data from the Atlantic Ocean, finding that PP at >10 μm has remained unchanged in most areas but exhibits an increasing trend in the South Subtropical Convergence and the Benguela Upwelling Zone. However, the present study is among the first to report long-term trends in PBc size fractionation in a freshwater ecosystem.

### Why do the PP and PBc L:S balances remain high?

There are three potential explanations for the PP and PBc L:S balances remaining high. First, the PBc of large phytoplankton may be inherently higher than that of small phytoplankton in coastal (Cermeño et al., 2005a; Maraóón et al., 2007) and upwelling (Cermeño et al., 2005b; Tilstone et al., 2017) areas. In the present study, *F*_*v*_*/F*_*m*_ and PB parameters of large phytoplankton were significantly elevated at the coastal site, i.e., Station 12C (Fig. 6). This suggests that coastal areas in both oceans and lakes are intrinsically conducive to large phytoplankton production, where nutrient limitation tends to be alleviated by mixing and terrestrial inputs. Indeed, TN and TP concentrations at Station 12C averaged 4.3 and 0.49 μmol L^−1^, respectively, over the 2020 study period, surpassing those at Station 12B (2.6 and 0.16 μmol L^−1^, respectively). Moreover, vertical mixing and light variability are favorable for large phytoplankton, which exhibit less sensitivity to light stress (Key et al., 2010; Kazama et al., 2022) as well as high light use efficiency (Finkel et al., 2010; Tilstone et al., 2017). These large phytoplankton in the coastal areas of Lake Biwa may be transported offshore by the first and second gyre, contributing to the maintenance of the population in the limnetic zone (Ishikawa et al., 2002).

The second possibility is the impact of increased solar radiation and transparency since the early 2000s (Fig. 1B, I). Indeed, a positive correlation was observed between the L:S balance of PP in 2020–2021 and the amount of solar radiation (Fig. 7). Large phytoplankton are adapted to intense light stress, potentially mitigating the negative effects of warming and oligotrophication. Yano et al. (2023) suggested that changes in dominant diatom species from *Skeletonema* (more susceptible to light stress) to *Chaetoceros* (less susceptible to light stress) in the Seto Inland Sea resulted from increasing solar radiation and oligotrophication. However, the extent of the increase in photon flux in the water column over these 30 years is unknown, and some data indicate that the photosynthetically active radiation in the surface layer has not significantly changed since the 2000s (Hayakawa, unpublished). More data are need to determine the impact of increased solar radiation on long-term PP and PBc L:S balances.

The third possibility is top-down effects due to zooplankton grazing. Previous studies have suggested that warming enhances zooplankton grazing, increasing the contribution of large phytoplankton to PP in mesocosms (Yvon-Durocher et al., 2015; Padfield et al., 2018). Indeed, zooplankton biomass has increased in recent years in Lake Biwa, attributed to the decrease in planktivorous fish abundance (Liu et al., 2019). Despite this, given that the seston C L:S balance showed a decreasing trend in 2020–2021, the zooplankton grazing effect on small phytoplankton may be negligible. Furthermore, the magnitude of warming may not be sufficient to increase zooplankton grazing rates or nutrient demand (Laspoumaderes et al., 2022).

This study has certain limitations. First, the absence of size-fractionated PP data for 1992 to 2020– 2021 adds uncertainty into the decadal trends of PBc L:S balance. As shown in Fig. 5, the PBc L:S balance varies with the year. Continuous monitoring of PP and PBc in a qualitative manner using a unified method is imperative for reliable assessments and evaluations of lake ecosystems. Second, caution should be exercised when interpreting the dynamics of PBc L:S balance as methodological differences exist between the present study and that of Urabe et al. (1995). Although we anticipate minimal discrepancies between gross and net PBc in slowly growing cells under oligotrophic conditions (Halsey & Jones, 2015), it is essential to recognize that the PBc L:S balance in 1992 (leaning toward *net* PBc) may not reflect the true *gross* PBc L:S balance. The relationship between gross and net PBc is affected by various factors, including carbon pool lifetimes and relative growth rates (Milligan et al., 2015), although the size dependency of these factors remains unclear in natural ecosystems. Third, building on the second point, a higher *gross* PBc L:S balance does not necessarily equate to a higher *net* PBc L:S balance. Although large cells can store substantial amounts of C, N, and P, their growth rate tends to be limited by the costs of material transport and assimilation rates (Marañón et al., 2013). Therefore, a qualitative assessment of net PBc is necessary to understand material cycling in a lake under the effects of climate change (Van de Waal & Litchman, 2020).

## Conclusions

Previous studies have predominantly focused on the relationships between climate change and/or oligotrophication and phytoplankton productivity solely through changes in biomass; thus, such studies lack the capacity to determine the temporal dynamics of productivity quantity and quality. The present study reveals that climate change and anthropogenic oligotrophication over the past 50 years have led to a decrease in PP in Lake Biwa since the 1990s, alongside changes in the phenology of PP dynamics. Furthermore, qualitative changes in PBc suggest that the PBc of large phytoplankton remains higher than that of small phytoplankton, with minimal changes observed since 1992. Three potential reasons for sustained large phytoplankton productivity include the favorable coastal environment, light conditions, and zooplankton grazing. Additionally, winter vertical mixing can sustain large phytoplankton populations in the upper surface layer (Hodoki et al., 2020). This study suggests that, even in a deep and oligotrophic lake such as in Lake Biwa, the production of large inedible phytoplankton does not decrease due to warming and oligotrophication. Rather, as total PP decreases while the PP L:S balance remains stable, the production of small edible phytoplankton decreases proportionally, potentially dampening trophic transfer efficiency and material cycling.

## Acknowledgements

We would like to thank Ayato Kozu and Hirokazu Teraishi for assisting with the chemical analysis. TK is an affiliated scientist at the Center for Ecological Research, Kyoto University.

## Supporting information

Appendix Table 1 and 2; Appendix Figs. 1–4

## Abbreviations

*F*_*v*_*/F*_*m*_: maximum PSII photochemical efficiency in the dark-adapted state
σ _PSII_(BGO): PSII light absorption cross-section under blue, green, and orange excitation flashes
PB_C_: carbon fixation rate per seston C per day
PB_C max_: maximum carbon fixation rate per seston C per day
α_C_: initial slope of the light-production curve based on seston C
PP: gross primary production per area per day

