## Appendix for "Impact of climate change and oligotrophication on quality and quantity of lake primary production: A case study in Lake Biwa"

Appendix Table 1 Steel's test for seston C, PP, and PBc L:S balance between 1992 and 2020–2021.

| Comparison | Estimator | Lower | Upper | Statistic | p.Value |
| --- | --- | --- | --- | --- | --- |
| Seston C L:S |  |  |  |  |  |
| p( 1992S , 2020 12BS ) | 0.47 | -0.34 | 1.27 | -0.17 | 1.00 |
| p( 1992S , 2020 12CS ) | 0.25 | -0.75 | 1.25 | -1.00 | 0.78 |
| p( 1992S , 2021 12BS ) | 0.30 | -0.42 | 1.02 | -1.11 | 0.72 |
| p( 1992S , 2021 12CS ) | 0.07 | -0.25 | 0.38 | -5.54 | 0.02 |
| PP L:S |  |  |  |  |  |
| p( 1992S , 2020 12BS ) | 0.33 | -0.28 | 0.94 | -0.90 | 0.79 |
| p( 1992S , 2020 12CS ) | 0.40 | -0.33 | 1.13 | -0.45 | 0.97 |
| p( 1992S , 2021 12BS ) | 0.43 | -0.21 | 1.08 | -0.34 | 0.99 |
| p( 1992S , 2021 12CS ) | 0.23 | -0.28 | 0.75 | -1.70 | 0.35 |
| PBc L:S |  |  |  |  |  |
| p( 1992S , 2020 12BS ) | 0.33 | -0.33 | 1.00 | -0.90 | 0.82 |
| p( 1992S , 2020 12CS ) | 0.50 | -0.35 | 1.35 | 0.00 | 1.00 |
| p( 1992S , 2021 12BS ) | 0.53 | -0.24 | 1.31 | 0.16 | 1.00 |
| p( 1992S , 2021 12CS ) | 0.53 | -0.19 | 1.26 | 0.17 | 1.00 |

Appendix Table 2 Steel's test for seston C, PP, and PBc between 1992 and 2020–2021.

| Comparison | Estimator | Lower | Upper | Statistic | p.Value |
| --- | --- | --- | --- | --- | --- |
| Seston C |  |  |  |  |  |
| p( 1992L , 2020 12BL ) | 0.22 | -0.31 | 0.75 | -1.77 | 0.36 |
| p( 1992L , 2020 12CL ) | 0.27 | -0.35 | 0.89 | -1.27 | 0.62 |
| p( 1992L , 2021 12BL ) | 0.19 | -0.30 | 0.69 | -2.06 | 0.25 |
| p( 1992L , 2021 12CL ) | 0.50 | -0.13 | 1.13 | 0.00 | 1.00 |
| p( 1992S , 2020 12BS ) | 0.22 | -0.31 | 0.75 | -1.77 | 0.36 |
| p( 1992S , 2020 12CS ) | 0.27 | -0.35 | 0.89 | -1.27 | 0.62 |
| p( 1992S , 2021 12BS ) | 0.19 | -0.30 | 0.69 | -2.06 | 0.25 |
| p( 1992S , 2021 12CS ) | 0.50 | -0.13 | 1.13 | 0.00 | 1.00 |
| PP |  |  |  |  |  |
| p( 1992L , 2020 12BL ) | 0.14 | -0.20 | 0.48 | -3.21 | 0.04 |
| p( 1992L , 2020 12CL ) | 0.00 | 0.00 | 0.00 | -849.77 | <0.001 |
| p( 1992L , 2021 12BL ) | 0.00 | 0.00 | 0.00 | -849.77 | <0.001 |
| p( 1992L , 2021 12CL ) | 0.03 | -0.09 | 0.15 | -12.02 | <0.001 |
| p( 1992S , 2020 12BS ) | 0.23 | -0.36 | 0.82 | -1.53 | 0.40 |
| p( 1992S , 2020 12CS ) | 0.00 | 0.00 | 0.00 | -819.94 | <0.001 |
| p( 1992S , 2021 12BS ) | 0.13 | -0.34 | 0.61 | -2.62 | 0.11 |
| p( 1992S , 2021 12CS ) | 0.10 | -0.27 | 0.47 | -3.65 | 0.04 |
| PBc |  |  |  |  |  |
| p( 1992L , 2020 12BL ) | 0.39 | -0.19 | 0.97 | -0.60 | 0.93 |
| p( 1992L , 2020 12CL ) | 0.60 | -0.05 | 1.25 | 0.48 | 0.97 |
| p( 1992L , 2021 12BL ) | 0.14 | -0.24 | 0.52 | -3.00 | 0.06 |
| p( 1992L , 2021 12CL ) | 0.28 | -0.24 | 0.79 | -1.36 | 0.51 |
| p( 1992S , 2020 12BS ) | 0.43 | -0.33 | 1.20 | -0.32 | 0.98 |
| p( 1992S , 2020 12CS ) | 0.55 | -0.32 | 1.42 | 0.21 | 1.00 |
| p( 1992S , 2021 12BS ) | 0.13 | -0.30 | 0.56 | -3.11 | 0.08 |
| p( 1992S , 2021 12CS ) | 0.30 | -0.45 | 1.05 | -0.98 | 0.68 |

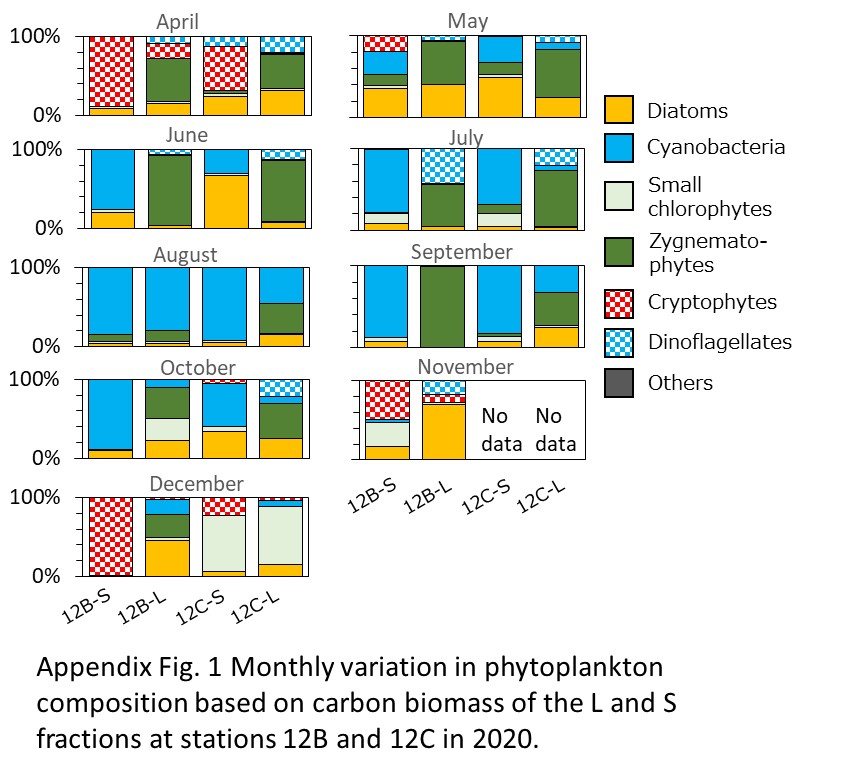

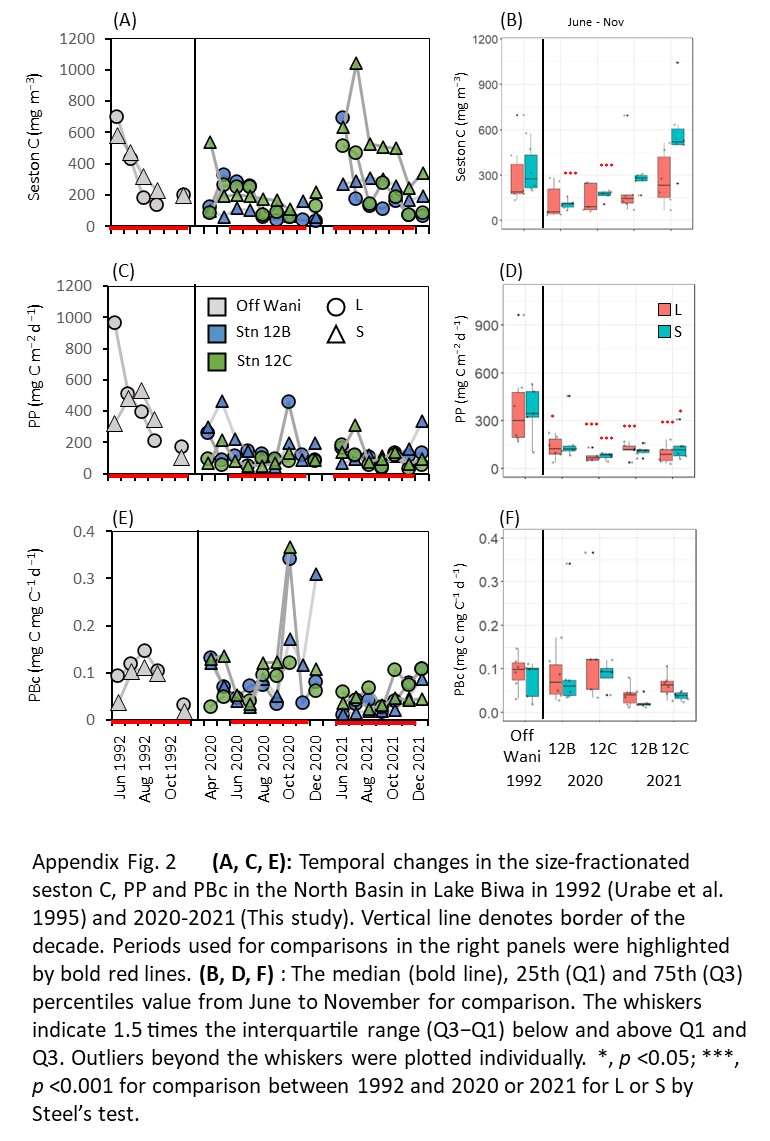

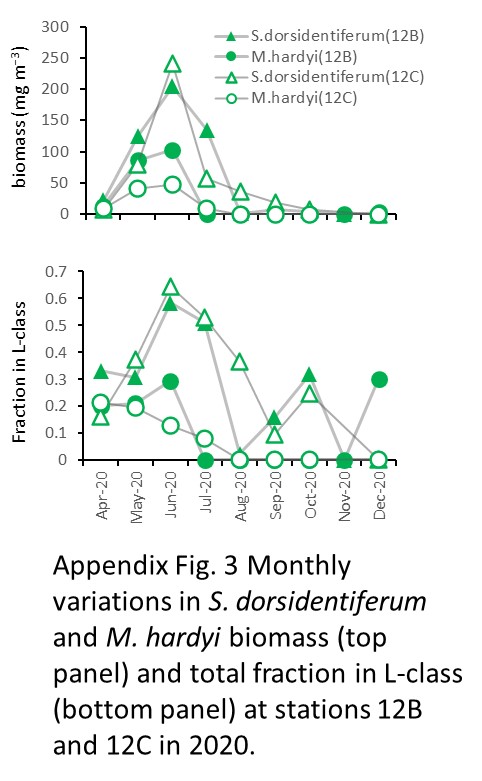

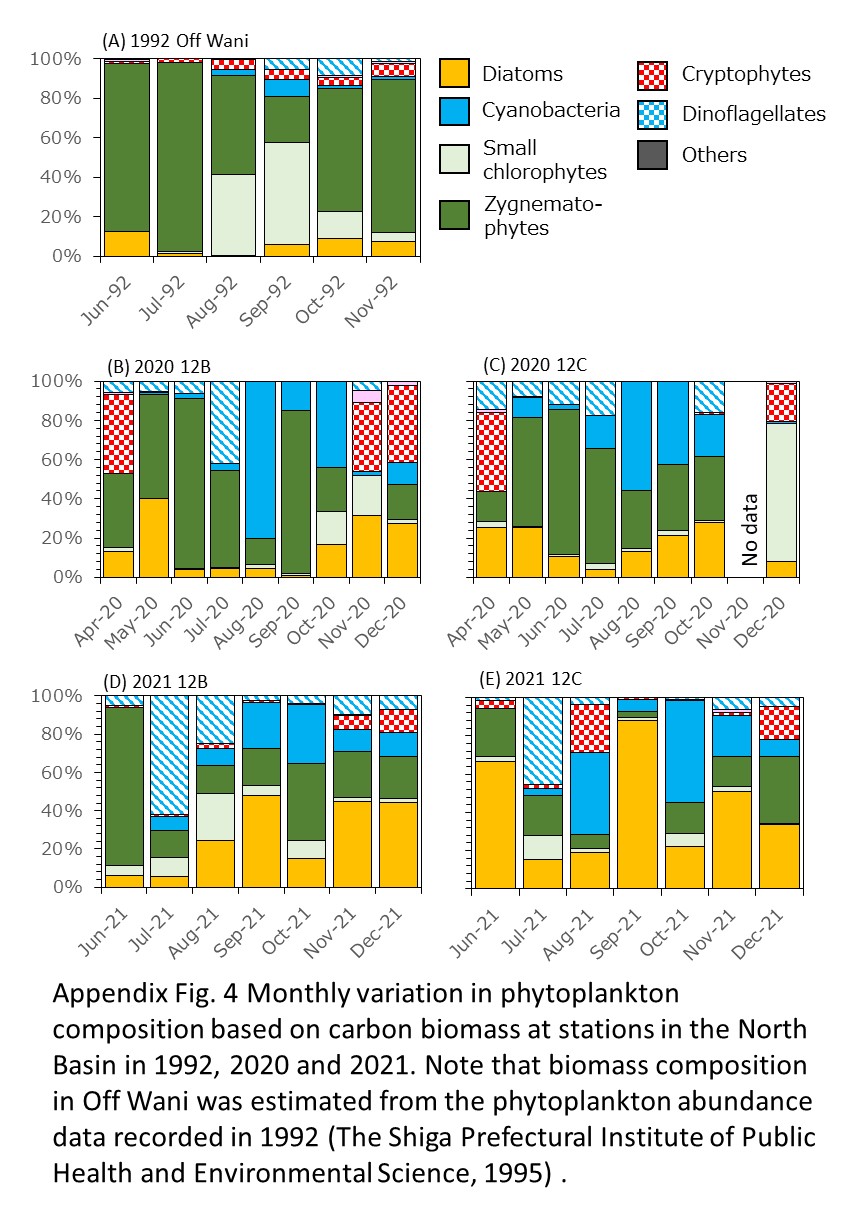
